# Impaired top-down auditory processing despite extensive single-neuron responses during human sleep

**DOI:** 10.1101/2021.04.03.438283

**Authors:** Hanna Hayat, Amit Marmelshtein, Aaron J. Krom, Yaniv Sela, Ariel Tankus, Ido Strauss, Firas Fahoum, Itzhak Fried, Yuval Nir

## Abstract

Sleep in all species is universally defined as a reversible, homeostatically-regulated state of a reduced behavioral responsiveness, with a high arousal threshold in response to external sensory stimulation^1^. However, it remains unclear whether sleep mainly gates motor output or affects responses along sensory pathways, and whether sleep primarily modulates specific aspects of the sensory response such as feedforward vs. feedback signaling^2–7^. Here, we simultaneously recorded polysomnography, iEEG, microwire LFPs, and neuronal spiking activity during wakefulness and sleep in 13 epilepsy patients implanted with clinical depth electrodes, while presenting auditory stimuli (e.g. click-trains, words, music). The results revealed robust spiking and induced LFP high-gamma (80-200Hz) power responses during both NREM and REM sleep across the lateral temporal lobe. The magnitude of the responses was only moderately attenuated in sleep, most notably for late responses beyond the early auditory cortex. Nonetheless, sleep responses maintained their tight locking with soundwave envelopes and their information content was only minimally affected. In contrast, a decrease in LFP alpha-beta (10-30Hz) power responses was prevalent in wakefulness but significantly disrupted in sleep. Entrainment to 40 Hz click-trains was comparable across REM sleep and wakefulness, but reduced in NREM sleep. In conclusion, our results establish the presence of extensive and robust auditory responses during sleep while LFP alpha-beta power decrease, likely reflecting top-down processes^8–10^, is deficient. More broadly, our findings suggest that feedback signalling is key to conscious sensory processing^11–13^.

## MAIN

The extent to which sleep affects responses along sensory pathways remains unclear. On one hand, external stimuli in sleep rarely affect perception^14^, suggesting response attenuation in cortical sensory regions. However, discriminative processing during sleep does persist for behaviorally-relevant or semantic incongruent stimuli^6,15–17^ as well as for contextual cues in targeted memory reactivation (TMR)^18,19^. Recent animal studies reporting comparable responses in the primary auditory cortex (A1) across sleep and wakefulness now challenge the long held assumption that natural sleep, like deep anesthesia, limits an effective relay to sensory cortex (‘thalamic gating)^20,21,2,3,22–24^. Whether this is also the case in consolidated human sleep remains unknown, since it is possible that robust auditory responses reflect a sentinel-like process that is unique to fragmented sleep in prey animals.

Previous non-invasive human studies using MEG ^5,25^, EEG^26–28^, and fMRI ^6,29^ have a number of limitations. Specifically, brief stimulation during sleep elicits a large stereotypical response (an evoked slow-wave often followed by a sleep spindle) known as a “K complex” that masks the precise dynamics and limits data interpretation. The spatial and temporal resolutions of EEG and fMRI, respectively, can not distinguish the neuronal sources of early (< 150 ms) selective auditory responses from late (~200-1000 ms) non-specific sleep responses^30^, and determine whether sleep predominantly affects feedforward or feedback processing.

To overcome previous limitations and compare auditory responses in wakefulness and natural sleep in humans, we recorded intracranial electroencephalogram (iEEG, n = 987 contacts), microwire local field potentials (LFPs, n = 937 microwires), and neuronal spiking activity (n = 713 clusters) from multiple cortical regions in 13 drug-resistant epilepsy patients implanted with depth electrodes for clinical monitoring (14 sessions, including eight full-night sessions lasting 484.8 ± 45.99 min, and six daytime nap sessions lasting 103.6 ±7.7min). At least one depth electrode in each monitored individual, targeted auditory (or other lateral temporal) cortical regions. We intermittently presented auditory stimuli including clicks, tones, chords, music, words, and sentences, via a bedside speaker (Methods) during the same recording session, while participants were awake or asleep (Fig. 1, Supplementary Table. 1). Sleep/wake stages were scored according to established guidelines^31^ (Fig. 1b, Extended Data Fig. 1) based on full polysomnography including electrooculogram (EOG), electromyogram (EMG), scalp EEG, and video monitoring whenever possible (n = 7 sessions), as previously described ^32^, or EEG/iEEG and video (n = 7 sessions, (Methods)).

**Figure 1.**
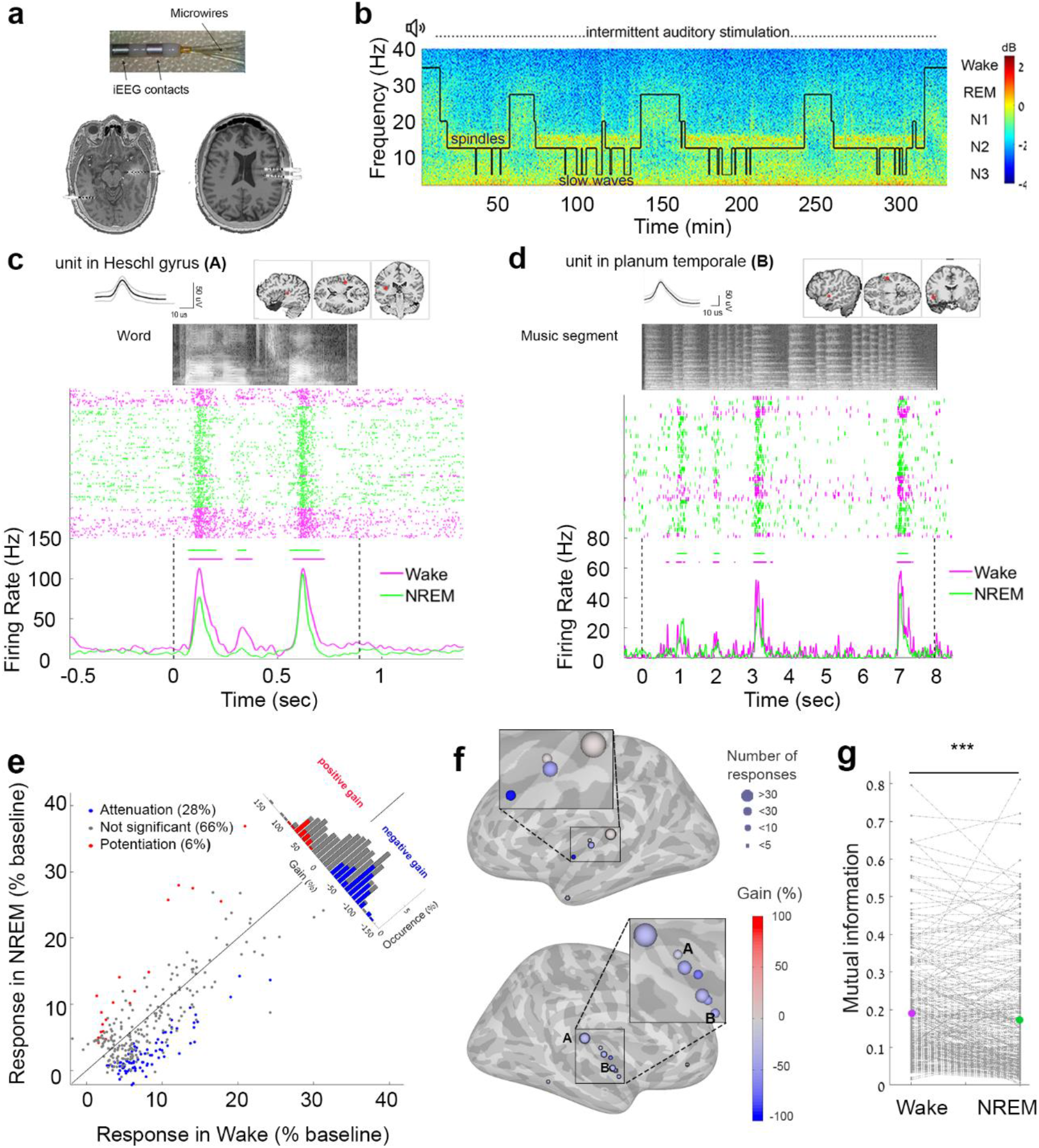
Robust auditory spiking responses across the temporal lobe during NREM sleep. (a) (top) Depth electrodes (6-12 per patient) implanted in epilepsy patients for clinical monitoring, each consisting of eight 1.5-mm iEEG contacts along the shaft and eight 40-μm microwires protruding from the distal tip, recording LFP and spiking activities, (bottom) two representative pre-implant MR images co-registered with post-implant CT used to localize electrode from the same individual. (b) Representative time–frequency representation (spectrogram) of iEEG recorded in one individual during a full-night sleep study with intermittent auditory stimulation. Warm colors (e.g. red, see color bar on far right) mark increased power in specific time–frequency windows (frequency shown on left side of y-axis). Superimposed hypnogram (gray trace) marks the time-course of sleep/wake states (shown on right side of y-axis). Note that NREM stages N2 and N3 are associated with increased power in spindle (10–15 Hz) and slow (< 4 Hz) frequency ranges. (c) Representative raster plots and peri-stimuli histograms (PSTHs) of unit spiking activities in response to auditory stimuli in the primary auditory cortex. (d) Same as (b) for a unit in higher-order auditory cortex (planum temporale). In both panels, the top row shows the action potential waveform (left inset, mean ± SD) and the anatomical location of the recorded unit (right inset, red circle in MRI sections), while the grayscale soundwave spectrograms are shown above the raster (lighter shades denote stronger power). Pink, wakefulness; Green, NREM sleep. Vertical dotted black lines mark stimulus onset and offset. Horizontal bars above PSTH time-courses indicate automatically-detected response intervals for which response magnitude was compared quantitatively. (e) Scatter plot of magnitude of all spiking responses to auditory stimuli (n = 312 responses from 55 clusters) in NREM sleep (y-axis) vs. wakefulness (x-axis), together with a histogram of gain values comparing response magnitudes (upper-right corner along the unity/zero-gain diagonal). (f) Gain values of spiking response magnitudes (NREM vs. wakefulness) in each region exhibiting auditory responses. The location of each circle denotes its anatomical location shown on a standard (MNI) brain template, the color represents the average gain detected in that region (color bar on right), and the circle’s size reflects the number of responses detected in that region. The letters A and B mark the location of representative units shown in panels A and B. (h) Mutual information (MI) between stimulus and spiking response (n = 312 responses from 55 neuronal clusters) in wakefulness (left, pink dot represents the mean) and NREM sleep (right, green dot represents the mean) reveals a modest (6.11%) reduction in MI during NREM sleep (***p < 0.001 by signed rank test).

We recorded spiking activity from 713 neuronal clusters (Supplementary Table 1), of which 55 clusters (7.7%) produced a significant auditory response (increased firing rate compared to baseline, p < 0.01 by Wilcoxon-Mann-Whitney test) to at least one stimulus in at least one vigilance state (n = 312 responses overall; Fig. 1c,d see Extended Data Fig. 2 for additional examples). The magnitude of most (66%) spiking responses was not significantly different between wakefulness and sleep, while the magnitude was significantly decreased in 28% and significantly increased in 6% (Fig. 1e, mean gain: −22.77%). The majority (293/312, 96%) of responses were observed in the superior temporal gyrus, but were also detected in other lateral temporal sites such as the middle temporal gyrus, as well as in the orbitofrontal cortex (Fig. 1f, Extended Data Fig. 3). Responses recorded in the posteromedial Heschl’s gyrus, probably corresponding functionally to A1^33,34^ (n = 236 in 33 neuronal clusters; mean gain = −16.45%), were significantly less attenuated than those in regions outside A1 (n = 91 in 22 neuronal clusters; gain = −37.58%, p = 0.00044 by Wilcoxon-Mann-Whitney test). However, most of the responses in both primary and non-primary regions, did not show significant attenuation (68% and 62% for A1 and outside A1, respectively). Indeed, even outside A1, robust high-fidelity responses persisted, as exemplified by the activity of a non-A1 neuronal cluster in response to presentation of an excerpt of Mozart music (Supplementary Video 1). Computing mutual information (MI) between the auditory stimulus and the spiking response (Methods) revealed a very modest decrease (6.11%, Fig.1g) in MI during NREM sleep compared to wakefulness, although this was nevertheless highly significant statistically (p < 10^−06^ by Wilcoxon signed rank test).

Since modulations in high-gamma (HG, 80-200 Hz) power are closely linked to neuronal firing rates in human auditory cortex^35^, we compared auditory-induced power responses across wakefulness and NREM sleep. The results revealed highly robust auditory-induced HG responses (Fig. 2a,b; additional examples in Extended Data Fig. 4; n = 554 responses from 74 LFP microwires and 311 responses in 47 iEEG electrodes). The majority (73%) of HG responses did not significantly differ in magnitude, although 18% were significantly attenuated in NREM sleep, and 9% were significantly potentiated (mean gain: −1.78%, Fig. 2c). In contrast to the spike responses, HG attenuation was higher in A1 (gain = −9.33%, n = 310 responses) than outside A1 (gain = 2.44%, n = 555 responses, p = 1.7*10^−05^ by Wilcoxon-Mann-Whitney test). The relationship of the gamma power envelope to the sound envelope of auditory stimuli was similar in sleep and wakefulness (Fig. 2e,f; mean correlation coefficients in wakefulness and sleep: r = 0.63 vs. r = 0.61, n = 424 responses in 41 LFP microwires, p = 0.72 by signed-ranked test). Auditory-induced power responses in the low-gamma (40-80Hz) range were significantly less attenuated in sleep than HG responses and displayed an average potentiation (Extended Data Fig. 5a, mean gain: +7.87%). There was a correlation between the sleep gain in each microwire and the gain of spiking responses in neuronal clusters identified on the same microwire (n = 45 LFP channels, r = 0.32 and r = 0.39 p=0.016 and p=0.005 for low-gamma, and high-gamma,_respectively, by random permutations tests, Methods, Extended Data Fig 5c, d). A higher response latency was strongly correlated with stronger sleep attenuation (Fig. 2g, r = 0.8, p < 0.001 by permutation). In addition, late/sustained components of the auditory response (> 200 ms) were associated with stronger sleep attenuation than early (< 200 ms) response components (Extended Data Fig. 6b, c). Other factors such as slow wave and spindle activities were also associated with higher sleep attenuation (Extended Data Fig. 6a). Comparing the degree of entrainment to fast stimulus modulations as described previously ^33^, we found that 40 Hz click trains in wakefulness strongly entrained field potentials (Fig. 2h, n = 281 iEEG channels and n = 84 LFP microwires, Methods). During NREM sleep, 25% of the iEEG and LFP electrodes exhibited significant attenuation compared to wakefulness, indicating a partial attenuation in 40 Hz entrainment (mean gain: −23.33%, n = 365 electrodes, Fig. 2i).

**Figure 2.**
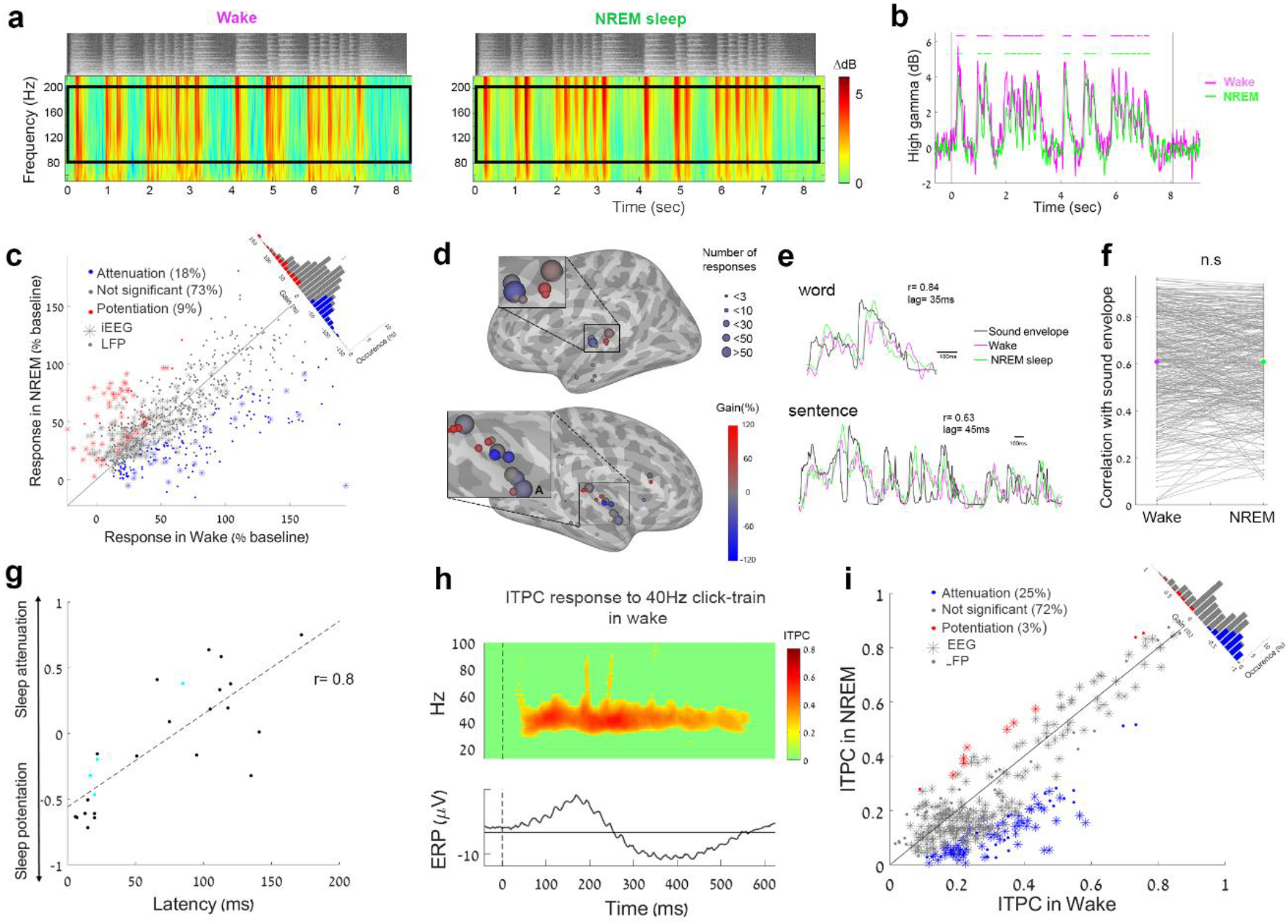
Robust induced high-gamma auditory responses across the temporal lobe and partially reduced ITPC during NREM sleep. **(a)** Representative spectrogram of induced LFP high-frequency (> 50Hz) power in response to music during wakefulness (left) and NREM sleep (right). Warmer colors (e.g. red) denote an increase in power (dB scale, color bar on right). Pink and gray rectangles represent time-frequency regions-of-interest used for subsequent quantification in wakefulness, and sleep, respectively. Grayscale soundwave spectrograms are shown above the LFP spectrograms (lighter shades denote stronger power). **(b)** Time-courses of the mean high gamma (80-200 Hz) responses shown in (A). Pink, wakefulness. Green, NREM sleep. Horizontal bars above the time-course indicate automatically-detected response intervals (Methods) for which the magnitude of the responses was compared quantitatively, Vertical black lines mark stimulus onset and offset. **(c)** Scatter plot of all high-gamma response magnitudes (% increase from baseline) to auditory stimuli (n = 556 responses from 74 LFP microwires and 320 responses from 55 iEEG channels) in NREM sleep (y-axis) vs. wakefulness (x-axis), together with a histogram of gain values comparing response magnitude (upper-right corner along the unity diagonal). **(d)** High gamma gain values (NREM vs. wakefulness) in each region exhibiting auditory gamma responses. The location of each circle denotes the anatomical location shown on a standard (MNI) brain template, while the color represents the average gain detected in that region (color bar on right), and the circle’s size reflects the number of responses detected in the region. The letter A marks the location of the representative microwire shown in panel A. **(e)** Two representative time-courses of LFP high-gamma responses (pink, wakefulness; green, NREM sleep) showing a tight relationship with the sound envelope of the auditory stimulus. Black, sound envelope; **(f)** Robust correlation between LFP high-gamma responses and the sound envelope is similar in wakefulness (left, pink dot represents the mean) and NREM sleep (right, green dot represents the mean; n = 424 responses in 41 microwires; p = 0.72 by signed rank test) **(g)** Scatter plot of the degree of response attenuation in NREM sleep (y-axis) vs. latency of gamma LFP response in each microwire (x-axis; n = 25 microwires, Pearson correlation coefficient r = 0.8, p < 0.001). Cyan dots mark individual adjacent microwires protruding from the same depth electrode exhibiting different sleep attenuations and latencies. **(h)** Top, iEEG ITPC (color map) in response to a 40 Hz click train. Bottom, event-related potential (ERP, black trace) of the same iEEG channel. **(i)** Scatter plot of inter-trial phase coherence (ITPC) of 40 Hz iEEG and LFP responses (n = 281 and 81, respectively) in NREM sleep (y-axis) vs. wakefulness (x-axis), together with a histogram of gain values comparing ITPC entrainment (along the diagonal).

In humans, sensory responses often manifest as an increase in spiking activity and LFP HG power, accompanied by a *decrease* in low-frequency power, also termed ‘desynchronization’ ^35–39^. Accordingly, during wakefulness we observed strong induced alpha-beta (10-30Hz) desynchronization (ABD), i.e. power suppression, in response to auditory stimuli (Fig. 3a,b; Extended Data Fig. 7). We proceeded to compare ABD responses (n=244 in 57 LPF microwires and n=188 in 29 iEEG electrodes) in wakefulness and NREM sleep. In sharp contrast to spiking and LFP HG responses, most (76%) ABD responses displayed significant attenuation during sleep, 23% did not change significantly, and only 1% were potentiated in sleep (Fig. 3c, mean gain: −71.84%). Similar results were obtained when considering each session as an independent variable (n = 8, mean gain: −56.97%, p = 0.004 by signed-rank test). Regional analysis (Fig. 3d) revealed that the ABD responses were less attenuated in A1 than outside A1 (mean gain: −41.44% vs. −80.20%; percent channels with significant ABD attenuation: 45% vs. 84% in A1 or outside A1, respectively, p = 7.3*10^−07^ by Wilcoxon-Mann-Whitney test). As observed for the HG responses, a longer latency was also associated with stronger ABD attenuation (Fig. 3e, r = 0.5, p < 0.001).

**Figure 3.**
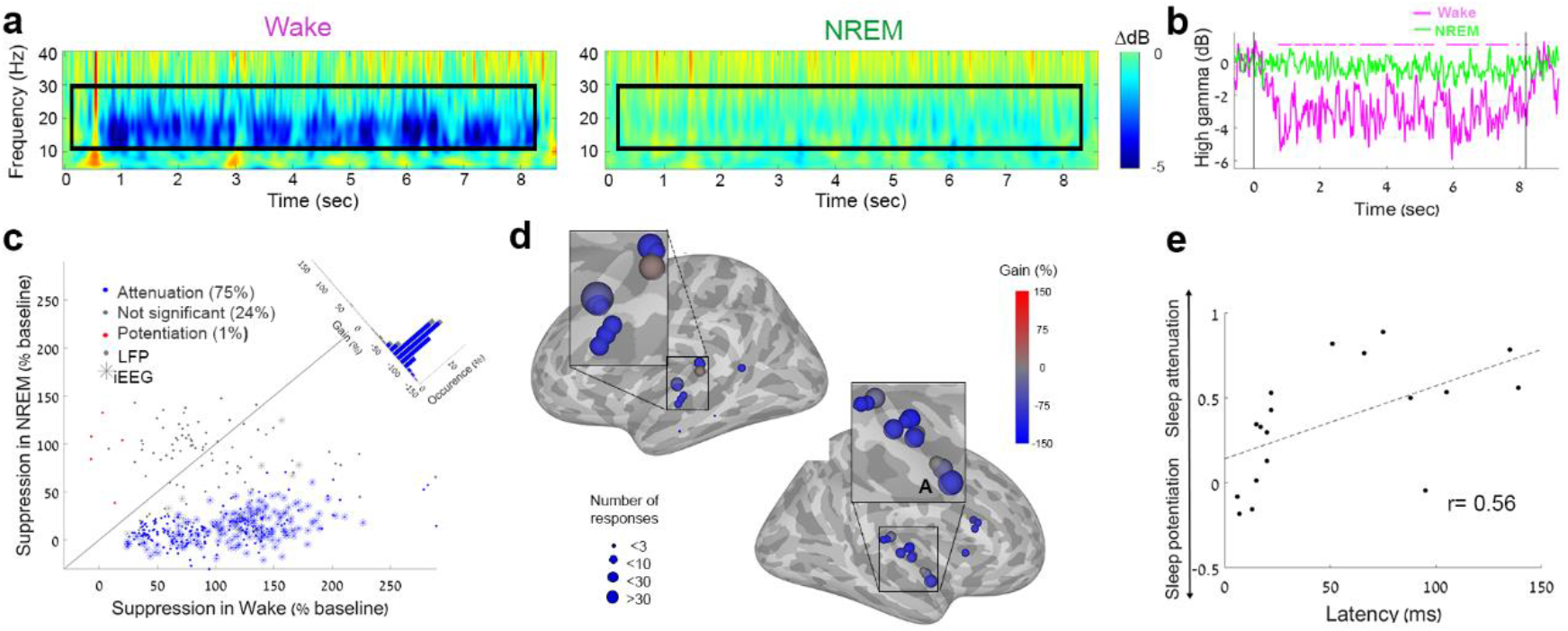
NREM sleep disrupts auditory-induced LFP alpha-beta power decreases. **(a)** Representative spectrogram of auditory-induced LFP low-frequency (< 50 Hz) power in response to music during wakefulness (left) and NREM sleep (right). Colder colors (e.g. blue) denote a decrease in power (dB scale, color bar on right). Pink and dark-green rectangles represent time-frequency regions-of-interest used for subsequent quantification in wakefulness, and sleep, respectively. **(b)** Time-course of induced alpha-beta (10-30 Hz) power dynamics shown in (a). Pink, wakefulness. Green, NREM sleep. Horizontal pink bars above the time-course indicate automatically-detected response intervals (Methods) for which the magnitude of the response was compared quantitatively (significant decreases were not detected in sleep), while vertical black lines mark stimulus onset and offset. **(c)** Scatter plot of all alpha-beta decrease (ABD) responses (% of baseline decrease) to auditory stimuli (n = 244 responses from 57 LPF channels and n = 188 responses from 29 iEEG channels) in NREM sleep (y-axis) vs. wakefulness (x-axis), together with a histogram of gain values comparing response magnitude (upper-right corner along the unity diagonal). **(d)** Differences in ABD responses (wakefulness vs. NREM) in each region exhibiting such responses. The location of each circle represents its anatomical location shown on a standard (MNI) brain template, the color reflects the average gain detected in that region (color bar on right), and the circle’s size reflects the number of responses detected in the region. The letter A marks the location of the representative microwire shown in panel A. **(e)** Scatter plot of the difference in ABD (wake-NREM, y-axis) vs. latency of ABD in each microwire (x-axis; n = 18 microwires, Pearson correlation coefficient r = 0.5, p < 0.001).

Lastly, we examined the auditory responses during REM sleep (n = 9 sessions in 8 patients). The results indicated that 85% of spiking responses (Fig. 4a, 141 responses from 25 neuronal clusters) did not significantly differ in magnitude between wakefulness and REM sleep, while the magnitude of 9% significantly decreased, and that of 6% significantly increased (Fig. 4b, mean gain: −16.75%). Similarly, the magnitude of induced HG responses was generally preserved during REM sleep (Fig. 4c, n = 236 LFP responses in 33 microwires; n = 198 iEEG responses in 33 electrodes), with 88% of responses with no significant changes, 7% of responses were significantly attenuated, and 5% were significantly potentiated (mean gain: −4.34%). ABD attenuation during REM sleep (n = 154 LFP responses in 32 microwires and n = 217 iEEG responses in 36 electrodes) showed a similar pattern to that found in NREM sleep: 46% responses were significantly attenuated, and 54% did not show any significant change (Fig. 4d, mean gain: −68.38%). There was significant correlation in the degree of attenuation during NREM and REM sleep in the same electrodes (r = 0.57 for HG responses and r = 0.51 for ABD, respectively, p < 0.001 by permutations, Extended Data Fig. 8 a,b). Notwithstanding the similarity between REM sleep and NREM sleep in terms of robust HG responses and significant ABD attenuation, entrainment to 40 Hz click-trains during REM sleep was not significantly different than that observed during wakefulness (Fig. 4d, mean gain: −3.72%, n = 242 iEEG electrodes and 60 LFP microwires, Extended Data Fig. 8c). Overall, during REM sleep, spiking responses displayed a modest attenuation, the profiles of the LFP/iEEG power responses were similar to those during NREM sleep, and 40 Hz entrainment was similar to that observed during wakefulness.

**Figure 4.**
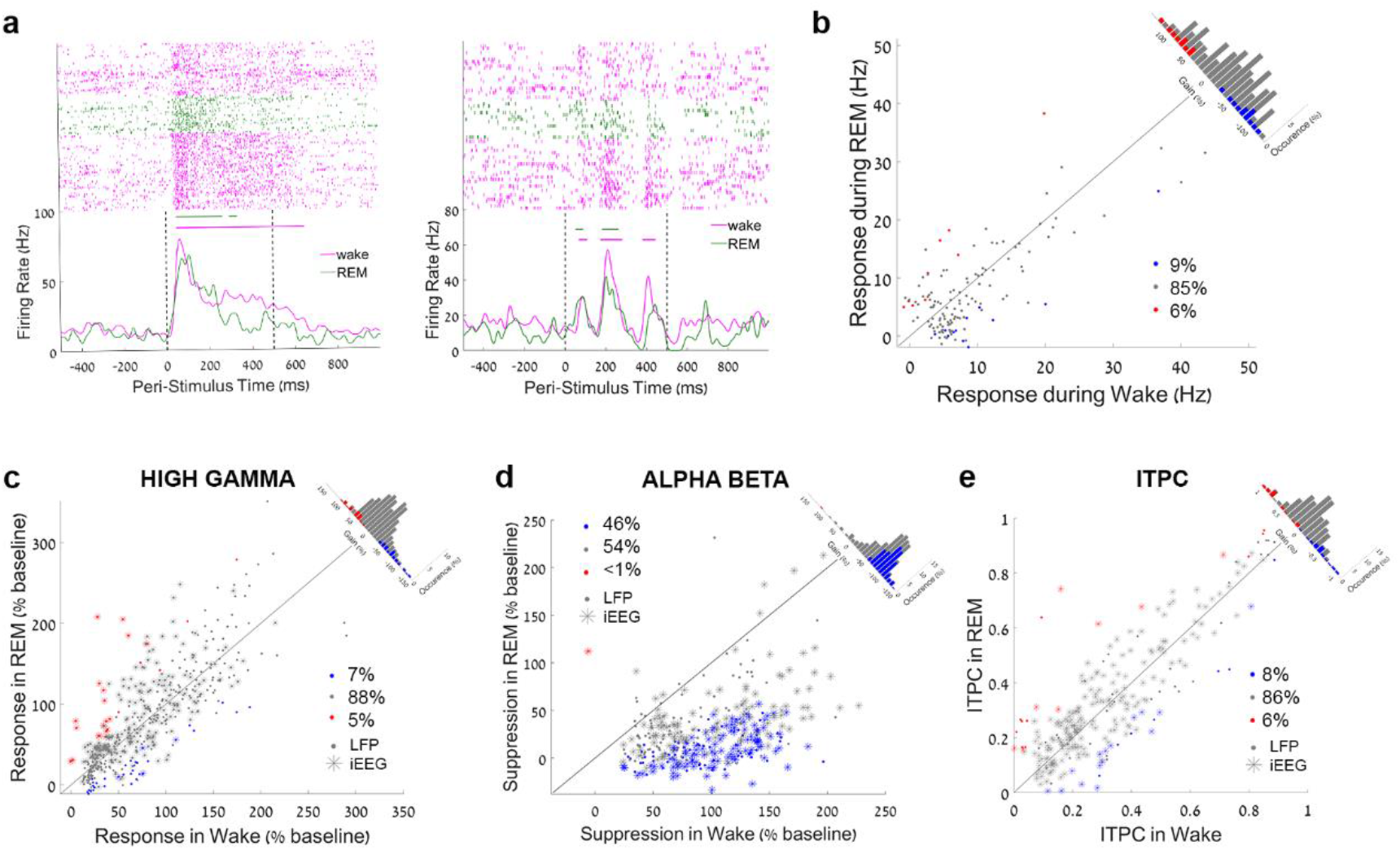
Auditory responses in REM sleep. **(a)** Two representative raster plots and peri-stimulus time histograms (PSTHs) of neuronal spiking activities in response to auditory stimuli (left: click train; right: word) in the primary auditory cortex. Pink, wakefulness; Green, REM sleep. Vertical dotted black lines mark stimulus onset and offset. Horizontal bars above the PSTH time-courses indicate automatically-detected response intervals (Methods) for which the magnitude of the response was compared quantitatively. **(b)** Scatter plot of all spiking responses to auditory stimuli (n = 141 responses from 25 clusters in REM sleep (y-axis) vs. wakefulness (x-axis), together with a histogram of gain values comparing response magnitude (upper-right corner along the unity/zero-gain diagonal). **(c)** Scatter plot of high-gamma responses (power increase) to auditory stimuli (n = 236 LFP responses from 33 channels and 198 iEEG responses from 30 channels) in REM sleep (y-axis) vs. wakefulness (x-axis), together with a histogram of gain values comparing the magnitude of the responses (upper-right corner along the unity diagonal, averaged gain of x %). **(d)** Scatter plot of all alpha-beta responses (power decrease) to auditory stimuli (n = 154 LFP responses from 32 channels and 217 iEEG responses from 36 channels) in REM sleep (y-axis) vs. wakefulness (x-axis), together with a histogram of gain values comparing the magnitude of the responses (upper-right corner along the unity diagonal, averaged gain of −71.7%). **(e)** Scatter plot of inter-trial phase coherence (ITPC) of 40 Hz iEEG and LFP responses (n = 174 iEEG electrodes and 28 LFP microwires) in REM sleep (y-axis) vs. wakefulness (x-axis), together with a histogram of gain values comparing ITPC entrainment (along the diagonal, averaged gain of x %) reveal comparable entrainment in the two vigilance states.

In summary, our results reveal robust neuronal and LFP HG auditory responses during sleep, well beyond the early auditory cortex. There was no significant attenuation of the magnitude of response for most responsive clusters/channels. Even the statistically significant attenuations were moderate in magnitude and more pronounced for late sustained responses and in regions downstream from A1. Responses during sleep continued to track the envelope of auditory soundwaves as they did in wakefulness, and their information content was only minimally reduced compared with that during wakefulness. In contrast to robust spiking/HG responses, LFP ABD was prevalent in wakefulness but significantly disrupted during both NREM and REM sleep. Finally, entrainment of field potentials to fast stimulus modulation rates (40 Hz click-trains) was reduced during NREM sleep, but comparable across desynchronized states of REM sleep and wakefulness. Our results establish that extensive and robust auditory responses persist during sleep while LFP alpha-beta power decrease is disrupted.

Some limitations inherent to research with neurosurgical epileptic patients should be acknowledged, but we do not believe that these play a major role or affect the conclusions. First, while we cannot entirely rule out the contribution of epileptiform activity, we carefully removed any epochs including signs of interictal epileptic activity from the analysis. The highly consistent results observed across patients with different clinical profiles argue against a major contribution by pathology. Second, the number of auditory responsive units is limited, but the significant correlation with LFP HG power modulations (Extended Data Fig. 7c and ^40^), allows us to extrapolate and confidently interpret the LFP HG responses as reflecting local spiking activities, even when limited in scope (e.g. in REM sleep). Third, we used a passive auditory stimulation paradigm. This approach could potentially limit the extent of responses (behavior- and task-related stimuli could drive responses in MTL mnemonic circuits^41^, not observed here), but importantly, this allowed us to address changes related to sleep *per-se*, without confounding processes such as reward or attention. Fourth, the localization of the electrodes did not permit a distinction between cortical layers.

Our results demonstrate the presence of robust neuronal and LFP HG power responses in the early auditory cortex with a similar magnitude of response in sleep and wakefulness. This is consistent with recent animal ^3,4,22,24^ and non-invasive human studies ^5,6,25,29^. There was a stronger attenuation during sleep in regions beyond the auditory cortex and in late sustained responses (Fig. 3g), as recently observed in the rat^4^. Notwithstanding the attenuation of responses downstream from A1 in some cases, most responses were not significantly decreased. In addition, spiking and HG exhibited high-fidelity responses as evidenced by mutual information analysis and tight locking to soundwave amplitude. Several lines of evidence suggest that the LFP gamma power responses are likely to represent feedforward (“bottom-up”) processing ^8,9,42,43^. Gamma oscillations are initiated in cortical input layer 4 and propagate to other cortical layers^8^. In addition, they are more readily observed in supragranular layers where feedforward projections originate^8,9,42,43^, they propagate from primary sensory regions to downstream high-level regions^8^, and blocking NMDA receptors and feedback processing boost their power^8^. We therefore interpret our results as representing a “feedforward sweep”^44^ in cortical sensory pathways that is tightly linked to physical stimulus features, but can not elicit sensory awareness on its own, as is the case in unconscious^45,46^ conditions such as anesthesia^33^.

Some aspects of the auditory response in REM sleep resemble those during NREM sleep, whereas other aspects are more similar to those in wakefulness, thus mirroring the general notion that REM sleep represents a ‘paradoxical’ hybrid of NREM sleep and wakefulness. LFP/iEEG induced power changes (HG increase, and ABD) were similar across sleep states (Fig. 4c,d; Extended Data Fig. 7), whereas time-locked entrainment to fast stimulus modulation rates (locking to 40 Hz click-trains) was lower in NREM sleep, but similar across wakefulness and REM sleep (Fig. 2i, Fig. 4e). Notably, NREM sleep and REM sleep share certain physiological aspects (e.g. low monoamine neuromodulation and low muscle tone^14^) and phenomenological aspects (e.g. disconnection from the external environment^14^). Accordingly, some aspects of altered sensory responses (e.g. robust LFP HG power changes with disrupted LFP ABD power changes) are also shared across NREM and REM sleep. Other physiological aspects of REM sleep more resemble those in wakefulness (e.g. high cholinergic tone, peripheral autonomic activation^14^) and the states also share certain phenomenological aspects (e.g. the ability to generate conscious experience). Accordingly, certain aspects of auditory responses, such as successful entrainment to fast stimulus modulation rates, are similar across REM sleep and wakefulness. Such entrainment is probably supported by desynchronized cortical activity enabled by high cholinergic tone^47^.

Our results point to alpha-beta desynchronization (ABD) as the most significant difference between sensory processing in wakefulness and sleep. ABD is readily observed in scalp EEG and intracranially upon auditory stimulation during wakefulness, even during passive listening ^39,48,49^, as well as in other brain regions and modalities^37,50^. Our results indicate that auditory-induced ABD during wakefulness is significantly disrupted during sleep (Fig. 3), as in anesthetic-loss of consciousness^33^. Under conditions examined to date, ABD exhibits high correlation with the degree of HG (although ABD is more spatially widespread) and the two phenomena can be parsimoniously described as a change in the exponent χ (“slope”) of the 1/f ^χ^ component of the spectrum^51^. However, we did not detect a significant correlation between the degrees to which sleep affected ABD and HG responses in individual electrodes, indicating that the two phenomena are largely independent^33,39^. A number of studies implicate ABD in top-down processing. In the macaque, gamma power propagates from V1 to V4 representing feedforward processing, whereas alpha oscillations propagate top-down from V4 to V1 mediating feedback processing^8^. Moreover, alpha (8–12 Hz) and beta (13–30 Hz) oscillations are maximal in infragranular layers^8,9,42^ where feedback connections arise^43^. ABD has also been shown to mediate feedback processing during visual stimulation^10,52^, is associated with better discrimination performance in the sensorimotor network^53^, and with the extent of auditory percepts in an illusory auditory paradigm^54^. The precise source of top-down signals remains elusive, and may include feedback from distant fronto-parietal regions, high-order sensory regions, or local recurrent networks in the early sensory cortex. Neuromodulatory systems are also likely to play a role, given their mediation of cortical desynchronization^47^, sensory perception^55^, and low activity in sleep^56^.

Thus, our study suggests that impaired top-down signaling is a key feature of sleep, and more generally, of loss of consciousness. Indeed, increasing evidence suggests that the neural mechanisms of anesthetic-induced unconsciousness may involve modulation of top-down processes ^57 58 59^. Anesthesia, and other unconscious states (e.g. vegetative states^11^), may decouple signaling along apical dendrites of layer 5 pyramidal neurons, thereby suppressing the influence of feedback arriving at the distal dendrites^13^. In conclusion, our results point to disrupted top-down signaling as a key feature of sleep, and to dissociation of feedforward and feedback signaling as a general feature of unconscious states and sensory disconnection.

## METHODS

### Patients

Thirteen drug-resistant epilepsy patients (five females) were implanted with Behnke-Fried depth electrodes (Ad-tech)^60^ as part of their clinical pre-surgical evaluation to identify seizure foci for potential surgical treatment. Electrode locations were based solely on clinical criteria. All patients provided written informed consent to participate in the research study, under the approval of the Institutional Review Board at the Tel Aviv Sourasky Medical Center (TASMC, 9 patients), or the Medical Institutional Review Board at UCLA (4 patients). In total, 14 sessions (6 naps / 8 nights) were recorded.

### Auditory stimulation

Auditory stimuli were delivered intermittently using a bed-side speaker during naps or full night sessions, where each recording session included periods of both wakefulness and sleep. Auditory stimuli were presented in a pseudo-random order, with the sound intensity level adjusted at the start of each session to be comfortably audible but not too loud, so the patients could sleep comfortably. Stimuli included 40 Hz click-trains, tone sequences, words, sentences, and music sequences [duration range: 0.5 s–9.4 s). Word stimuli were compiled in English at UCLA and in Hebrew at TASMC.

### Sleep staging

Full polysomnography (scalp EEG, EOG, EMG, and video) was recorded in seven sessions (three nights / four naps). Epochs were scored as wakefulness (W), N1/unknown, N2, N3, and REM sleep according to established guidelines ^31^. In three sessions (2 nights / 1 nap), only the scalp EEG signal was recorded together with intracranial data. In these cases, sleep scoring was performed using the scalp EEG, confirmed by visualization of iEEG spectrograms and video recordings. Periods scored as N2 and N3 displayed high levels of SWA and sigma (sleep spindle) activity, whereas periods of wakefulness and REM sleep were associated with low levels of SWA. For four sessions (two nights & two naps), sleep scoring was based on iEEGs and video recordings. We calculated time-frequency dynamics of the iEEG (spectrograms) using a 30 s window (without overlap) spanning frequencies from 0 to 40 Hz and averaged the power in the delta band (0.5-4 Hz). Epochs with delta power higher than the 55^th^ percentile were scored as NREM sleep and those with delta power lower than the 20^th^ percentile, were scored as wakefulness/REM sleep and were further subdivided as: epochs where the video showed that the patient was awake (eyes open, moving, sitting) were scored as wakefulness. Long periods (> 3min) occurring during the second part of the night, where the video indicated that the patient was likely to be asleep (closed eyes, no movements), were scored as REM sleep. To further validate sleep scoring based solely on iEEG, we compared our automatic sleep scoring to manual scoring in the overnight sessions with full polysomnography (PSG). The results indicated that 81.47 ± 12.36%, 88.33 ± 6.68%, and 84.44 ± 6.51% (n = 4 nights) of the epochs scored by automatic scoring as wake, NREM sleep and REM sleep, respectively, agreed with the scoring labels obtained by full PSG.

### Electrophysiology

Each depth electrode had eight platinum iEEG contacts along the shaft (referenced to the scalp), together with eight micro-wires protruding 3-5 mm from the distal tip, and a 9^th^ low-impedance reference microwire ^60^ that served as referenced for each microwire electrode bundle. Data were recorded using either Blackrock (Salt Lake City, UT, USA, 30 kHz sampling rate) or Neuralynx (Bozeman, MT, USA, 40 kHz sampling rate) data acquisition systems.

### Spike sorting

Neuronal clusters were identified using the ‘waveclus’ software package ^61^ as described previously^62,63^: extracellular recordings were high-pass filtered above 300 Hz and a threshold of 5 standard deviations above the median noise level was computed. Detected events were clustered (or categorized as noise) using automatic superparamagnetic clustering of wavelet coefficients, followed by manual refinement based on the consistency of spike waveforms and inter-spike interval distributions.

### Detection of significant responses

We identified neuronal auditory responses as described previously ^4^. First, the response in each trial was smoothed by convolution with a Gaussian kernel (σ = 10 ms). Next, a one tailed Wilcoxon-Mann-Whitney test was used to compare the results across trials. Each millisecond (within an interval corresponding to the stimulus duration + 100 ms following it) was compared against baseline activity (we corrected for the multiple comparisons using False-Discovery-Rate ^64^ with base alpha of 0.01). A minimum of six trials per condition (wakefulness or sleep states) was required. Components shorter than 5 ms were excluded, and undetected intervals shorter than 2 ms that preceded and followed responses were categorized as misses and bridged with adjacent intervals. To further reduce the risk of false detections, the total length of the response for each stimulus had to be greater than 1.5% of the stimulus length. Responses were normalized by subtracting the pre-stimulus baseline (0-500 ms) activity in each state (baseline normalization).

### Mutual Information Analysis

To estimate how informative the spiking response of each unit was with respect to the set of temporally dynamic stimuli (various words, click-trains, music segments, and tones), we divided each stimulus into 50 ms bins and calculated the number of spikes per bin for each trial and stimulus (e.g. a word of 450 ms duration was segmented to nine consecutive bins). We then pooled together the bins of all stimuli and calculated the mutual information between the two discrete variables of spikes count in each bin (*r*, response) and the bin identity (*s*, stimulus):

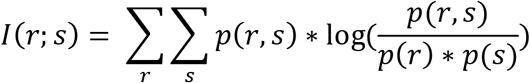

When comparing the mutual information between different behavioral states, the number of trials for each stimulus was equalized across states. Qualitatively similar results were obtained for 20, 50 and 100 ms bins, suggesting that the choice of a 50 ms bin size did not affect the results.

### LFP and iEEG power analysis

Signals from macro- and micro-electrodes were down-sampled to 1 kHz and band-pass filtered between 40-80 Hz, 80-200 Hz, and 10-30 Hz for low gamma, high gamma, and alpha-beta frequency bands, respectively. They were then Hilbert-transformed to obtain the instantaneous amplitude envelope, and log converted to express their amplitude in dB. For each channel and frequency band, the baseline power was extracted from a 500 ms interval before trial onset, and the mean baseline power was subtracted from the response power, separately for each frequency-band of interest and separately for each channel. Trials with power higher than 5 std from the mean were excluded.

Time intervals associated with significant induced LFP power in response to auditory stimuli were detected with the same method described above for the neuronal response. For LFP responses, response components shorter than 10 ms for low and high gamma (and 50 ms for alpha-beta) were excluded, and undetected intervals shorter than 4 ms that preceded and followed responses were categorized as misses and bridged with adjacent intervals. All responses were also inspected visually to rule out false automatic detections. These parameters were optimized after extensive visual inspection of automatic response detections; importantly, none of the results reported were dependent on the precise parameters used for response detection.

For latency analysis, the same automatic algorithm was applied on low-gamma filtered channels that exhibited a significant response to 40 Hz click-trains during the first 200 ms of the response interval. The first time point in this interval that showed significantly higher activity than baseline was defined as the response latency.

### Comparison across vigilance states (LFP analysis and spiking activity)

For each stimulus and pair of states to be compared (e.g. wakefulness vs. NREM sleep), we separately identified temporal intervals with significant responses in either state as described previously ^4^. A two tailed Wilcoxon-Mann-Whitney test was used to compare the response in wake and sleep (alpha of 0.01) for each stimulus.

We quantified the relation between response magnitudes in wakefulness and sleep using a gain factor as described previously ^3,4,24,33^, after normalizing each response to the baseline of that state:

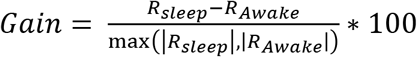 where RSleep and RAwake are the response amplitudes for a specific cluster/channel during wakefulness or sleep.

To compare response magnitudes between vigilance states per neuronal cluster (spikes) or channel (LFP/iEEG) (see Extended Data Fig. 5c,d), we combined the responses from all stimuli (same number of trials per stimulus), and then calculated the mean response for each channel/neuronal cluster in each state (wake/NREM sleep).

### Analysis of correlation with soundwave envelope

LFP microwires with gamma band power modulations that displayed a significant response, were further analyzed to quantify their correlation with soundwave envelope (intensity dynamics). The soundwave envelope was extracted by calculating the running average of the square amplitude using a 5 ms window (without overlap). The high gamma response was down-sampled to 200 Hz. We first identified, using cross-correlation, the temporal lag associated with the highest correlation. This was followed by calculation of the Pearson correlation between the response time-course and the soundwave envelope at this time lag, and analysis of the statistical significance using permutations (p < 0.01).

### Inter-trial phase coherence (ITPC) analysis of responses to 40 Hz click-trains

Responses to 40 Hz click-train were quantified using inter-trial phase coherence (ITPC), calculated as described previously ^33^. Briefly, ITPC was defined as: 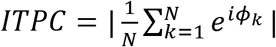 where N represents the number of trials and φk the phase of the spectral estimate for trial k for the 40 Hz frequency.

### Slow wave activity and sigma power analysis

For each session, we calculated the power spectrum of the scalp EEG in the 2 s interval preceding stimulus onset (or iEEG), and extracted the slow wave activity (SWA, 0.5-4 Hz) and the sigma power (10-16 Hz). For each stimulus eliciting a significant response, we sorted the trials according SWA and separated trials occurring during low SWA (below the 20th percentile) or during high SWA (above the 80th percentile). A minimum of 10 trials in each category was required to include a specific channel in this analysis. We then compared the response for each stimulus between the two groups by Mann-Whitney tests.

### Electrode localization

Pre-implant MRI scans (Siemens Prisma scanner or Magnetom Skyra or GE Signa scanner, 3T, T1 sequence, resolution 1 mm × 1 mm × 1 mm or 1 mm × 1 mm × 5 mm) were co-registered with post-implant computed tomography (CT) scans (Philips MX8000 or Brilliance or Siemens Sensation-64, resolution 1.5 mm × 0.5 mm × 0.5 mm or 0.75 mm × 0.5 mm × 0.5 mm) to identify the locations of the electrodes. Individual subject data were further transformed into brain average space to facilitate the simultaneous visualization of electrode positions in different individuals. Co-registration and localization were estimated by using FreeSurfer^65^ and BioImage^66^ software, according to the guidelines of iELVIS^67^.

## ACKNOWLEDGEMENTS

Supported by the Israel Science Foundation (ISF) grants 1326/15 (YN), 762/16 (AJK), and 51/11 (I-CORE cognitive sciences; YN), National Science Foundation & US-Israel Binational Science Foundation (NSF-BSF) grant 2017628 (IF and YN), the Adelis Foundation (YN), the European Research Council (ERC-2019-CoG 864353; YN), the European Society of Anesthesiology (AJK) and by an Azrieli Foundation fellowship award (Y.S). We thank Michelle Tran, Guldamla Kalender and Natalie Cherry for assistance at UCLA, Yitzhak Norman for iELSVIS training, Maya Geva-Sagiv for helpful comments and all Nir lab members for discussions.

## AUTHOR CONTRIBUTIONS

Y.N. and I.F. conceived research and secured funding. H.H., A.M. and A.J.K. designed experiments and collected data. H.H, A.M., A.J.K and Y.S analyzed data, supervised by Y.N.. I.F. and I.S. performed surgeries. F.F. supervised clinical care at TAMSC and analyzed epilepsy profiles. A.T. managed clinical electrophysiology setup. H.H., A.M., and Y.N. wrote the manuscript. All authors provided ongoing critical review of results and commented on the manuscript.

## DATA AVAILABILITY

Data sets supporting the findings of this paper are available on request from the corresponding authors.

## EXTENDED DATA AND FIGURES

**Supplementary Table 1.** Data acquisition details. Data acquisition details for the nine recording sessions included in this study. Patients d009 and d011 refer to the same individual who was re-admitted with new electrode locations due to inconclusive clinical results from the first hospital admission. Abbreviations: UCLA= University of California Los Angeles, TASMC=Tel Aviv Sourasky Medical Center, 40 Hz CT= 40 Hz click-train

| Patient | 472 | 479 | 505 | 489 | D009 | D011<br>2 sessions | D013 | D014 | D017 | D018 | D023 | D025 | D026 |
| --- | --- | --- | --- | --- | --- | --- | --- | --- | --- | --- | --- | --- | --- |
| Session | Nap | Nap | Nap | Nap | Nap | Night | Night | Night | Night | Nap | Night | Night | Night |
| Gender | F | M | M | F | F | F | M | M | M | M | W | M | M |
| Age | 43 | 35 | 35 | 47 | 24 | 25 | 35 | 28 | 39 | 29 | 34 | 17 | 27 |
| Location | UCLA | UCLA | UCLA | UCLA | TASMC | TASMC | TASMC | TASMC | TASMC | TASMC | TASMC | TASMC | TASMC |
| Seizure Onset Zone | right OF and left anterior temporal | Mostly temporal | left anterior fusiform gyrus and second connected to left oblique insula | Left mesial temporal lobe (LEC), with one seizure in the lateral RAH | Right temporal / frontal lobes | Right superior frontal gyrus | Right temporal tip | Left mesial frontal | zone unclear | Right frontal lobe | Right posterior mesial temporal | Right anterior superior temporal gyrus | Right orbito-frontal |
| Stimuli | Words, music, sentence 40 Hz CT | Words, music, sentence 40 Hz CT | Words, music, sentence 40 Hz CT | Words, music, sentence 40 Hz CT | Words, 40Hz CT | Words, music, sentence 40 Hz CT | Words, music, sentence 40 Hz CT | Words, music, sentence 40 Hz CT | Words, music, sentence 40 Hz CT | Words, music, sentence 40 Hz CT | Words, music, sentence 40 Hz CT | Words, music, sentence 40 Hz CT | Words, music, sentence 40 Hz CT |
| Language | English | English | English | English | Hebrew | Hebrew | Hebrew | Hebrew | Hebrew | Hebrew | Hebrew | Hebrew | Hebrew |
| Total N. of units | 74 | 150 | 102 |  | 62 | 45 | 0 | 0 | 55 | 60 | 83 | 58 | 24 |
| N. of responsive units | 7 | 25 | 14 | 5 | 1 | 2 | 0 | 0 | 1 | 0 | 0 | 0 | 0 |
| N. of responsive HG (LFP / iEEG) | 6 (0) | 24 (14) | 16 (0) | 14 (0) | 4 (5) | 10 (7) | 0 (1) | 0 (0) | 0 (0) | 0 (0) | 0 (10) | 0 (0) | 0 (0) |
| N. of responsive ABD (LFP / iEEG) | 4 (0) | 16 (10) | 12 (0) | 2 (0) | 7 (7) | 12 (11) | 0 (7) | 0 (0) | 0 (7) | 0 (0) | 0 (0) | 0 (0) | 0 (0) |
| N. of responsive ITPC to 40 Hz CT (LFP / iEEG) | 1 (2) | 57 (25) | 3 (10) | 24 (11) | 11 (1) | 39 (13) | 40 (5) | 1 (1) | 1 (2) | 27 (0) | 47 (10) | 26 (2) | 4 (2) |

**Extended Data Fig. 1.**
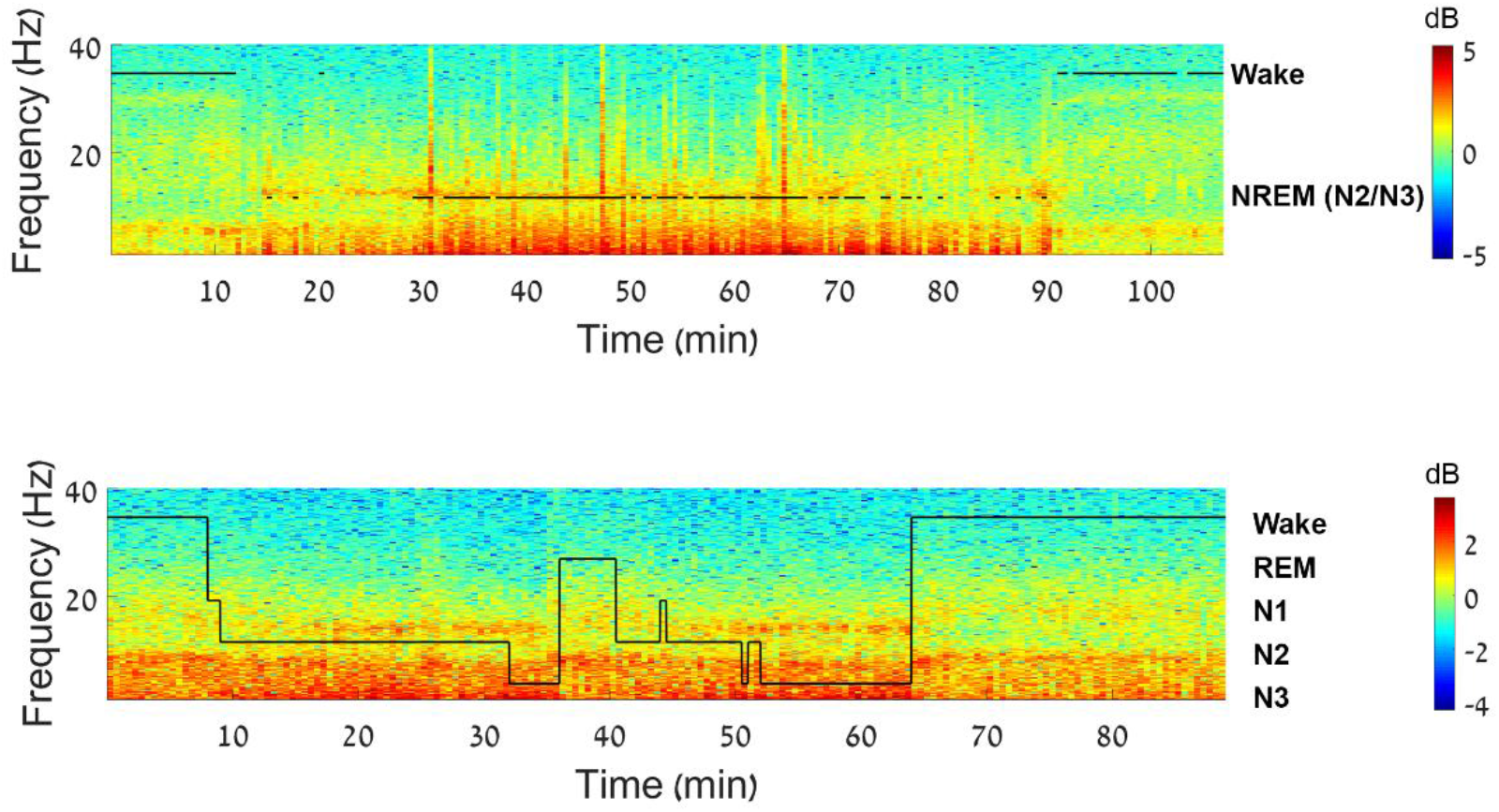
Sleep scoring. Representative time–frequency representation (spectrogram) of iEEG recorded during a nap session. Warm colors (e.g. red) indicate increased power in specific time–frequency windows (frequency shown on left side of y-axis). Superimposed hypnograms (in black) present the time-course of sleep/wake states (shown on right side of y-axis); top, one nap session with automatic sleep scoring; and bottom, one nap session with full PSG.

**Extended Data Fig. 2.**
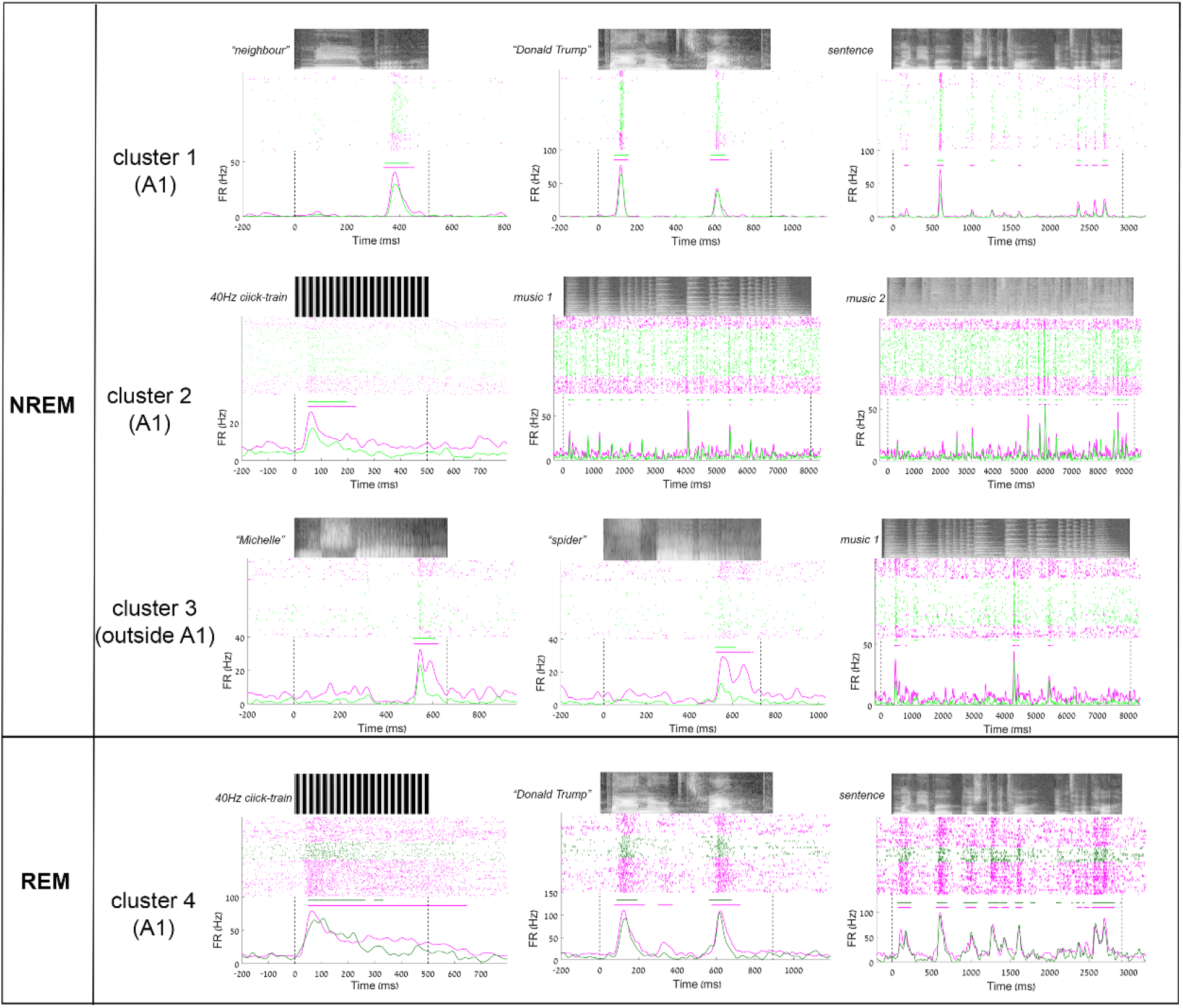
Additional examples of neuronal auditory responses during wakefulness and NREM sleep. Representative raster plots and PSTHs of unit spiking activities in auditory cortex in response to auditory stimuli during wakefulness (pink) and NREM sleep (light green) or REM sleep (dark green). Grayscale soundwave spectrograms are shown above each raster (lighter shades denote stronger power). Vertical dotted black lines mark stimulus onset and offset. Horizontal bars above PSTH time-courses indicate automatically-detected response intervals (Methods) for which the response magnitudes were compared quantitatively.

**Extended Data Fig. 3.**
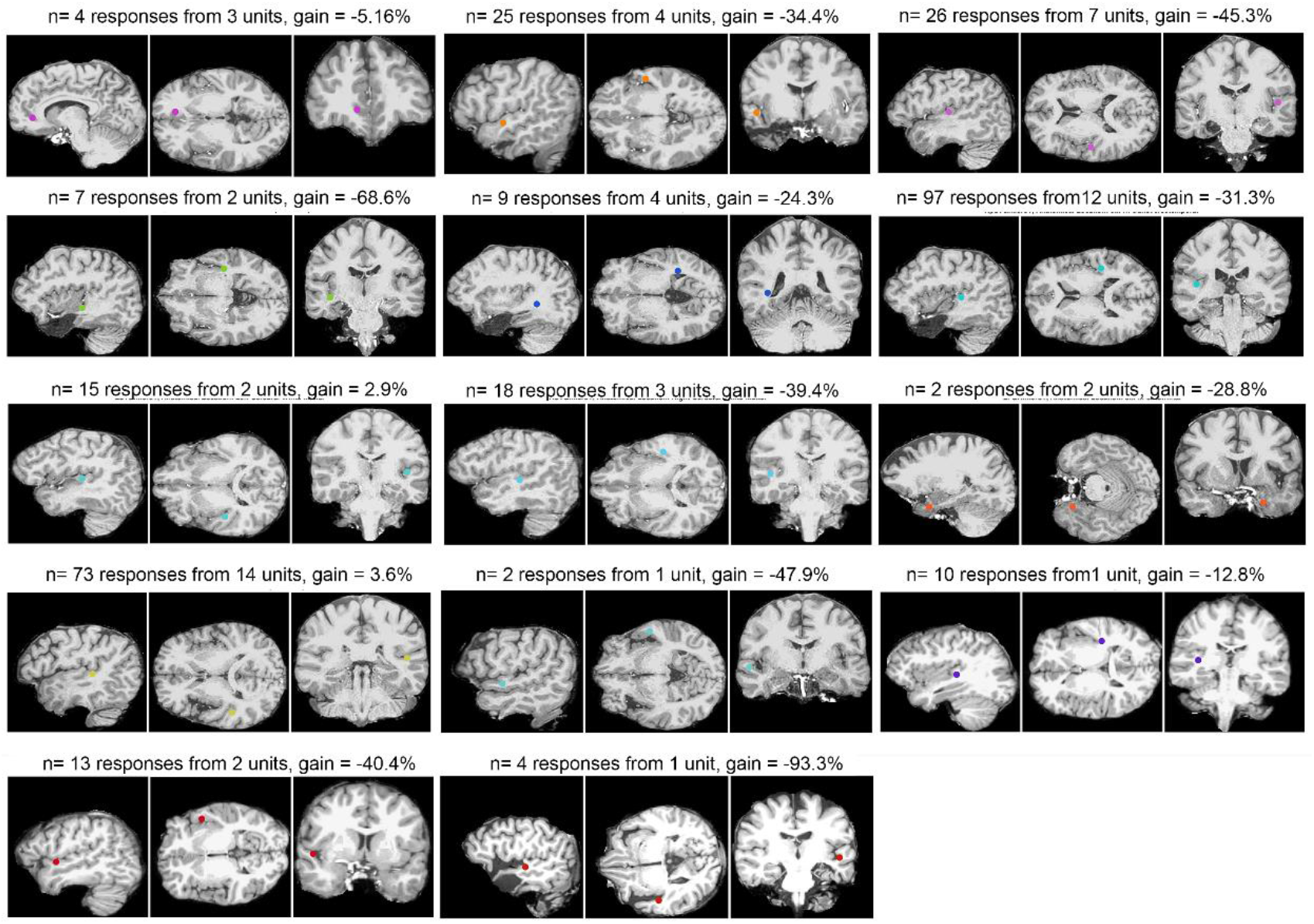
Anatomical location of auditory-responsive units. Each triplet of brain images shows sagittal (left), axial (middle), and coronal (right) MR sections. Colored dots denote the location of the microwire bundle as identified by co-registration of post-implant CT with pre-implant MRI (Methods), using native (individual) patient coordinates

**Extended Data Fig. 4.**
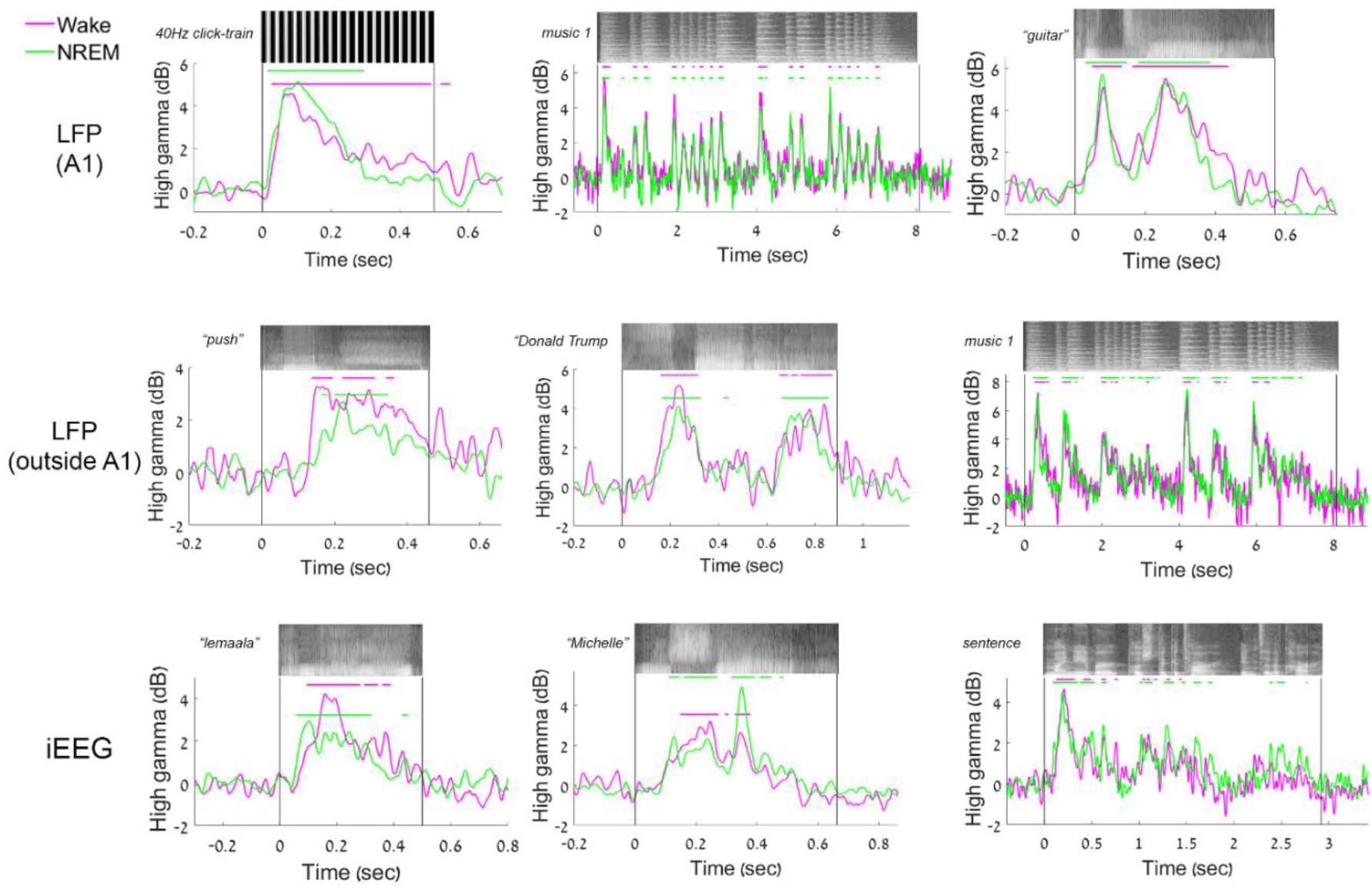
Additional examples of LFP induced high-gamma auditory responses during wakefulness and NREM sleep. LFP induced high-gamma (80-200 Hz) power time-courses during wakefulness (pink) and NREM sleep (green) reveal similar profiles.

**Extended Data Fig. 5.**
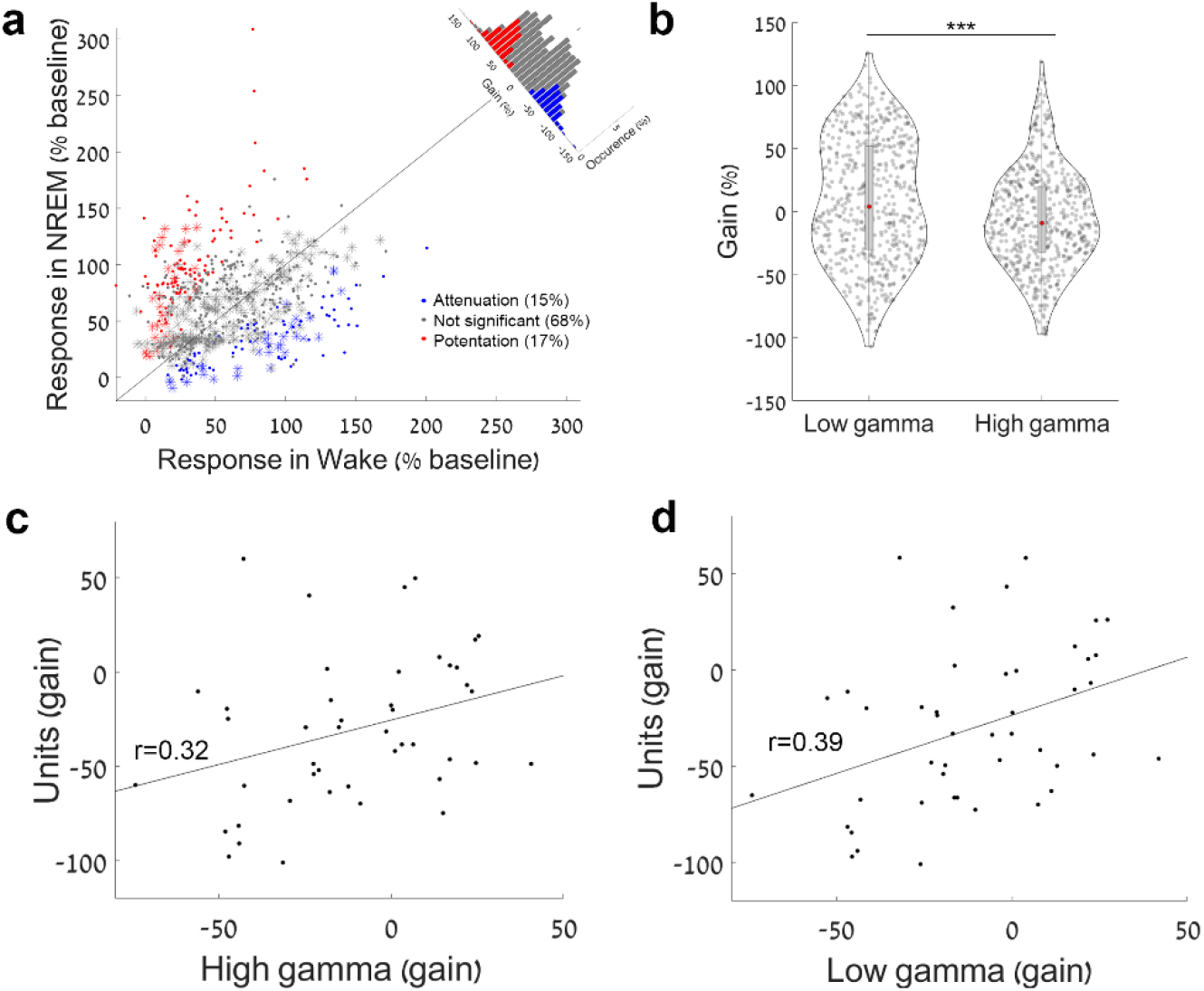
Comparing how NREM sleep affects responses in spiking activity, LFP low-gamma power, and LFP high-gamma power. (a) Scatter plot of all low-gamma (40-80 Hz) responses (power increase) to auditory stimuli (n = 418 responses from 61 LFP microwires and 292 responses from 42 iEEG electrodes) in NREM sleep (y-axis) vs. wakefulness (x-axis), together with a histogram of gain values comparing response magnitude (upper-right corner along the unity diagonal). Sleep was not associated with a trend for attenuation in response magnitude; instead, most (68.03%) low-gamma responses did not show significant differences across states, with14.51% significantly attenuated in NREM sleep, and 17.46% significantly potentiated in sleep (mean gain: +7.87%). (b) Scatter plot of NREM sleep attenuation in LFP high-gamma responses (y-axis) vs. NREM sleep attenuation in LFP low-gamma responses (x-axis) reveals stronger sleep attenuation in high-gamma responses for the same microwires and stimuli (556 responses from 87 channels, mean gain: +8.7% and −5.6% for low gamma and high gamma, respectively, p < 0.001 by Wilcoxon signed-rank test. (c) Scatter plot of the degree of NREM sleep attenuation in spiking responses (y-axis) vs. the degree of NREM sleep attenuation in LFP high-gamma responses (x-axis) reveals significant correlation (n = 45 channels/units respectively, Pearson correlation coefficient r = 0.32, p = 0.016, by random permutations tests, (Methods). (d) Same as (c) for LFP low-gamma power (Pearson correlation coefficient r = 0.39, p = 0.005, n = 40 channels/units).

**Extended Data Fig. 6.**
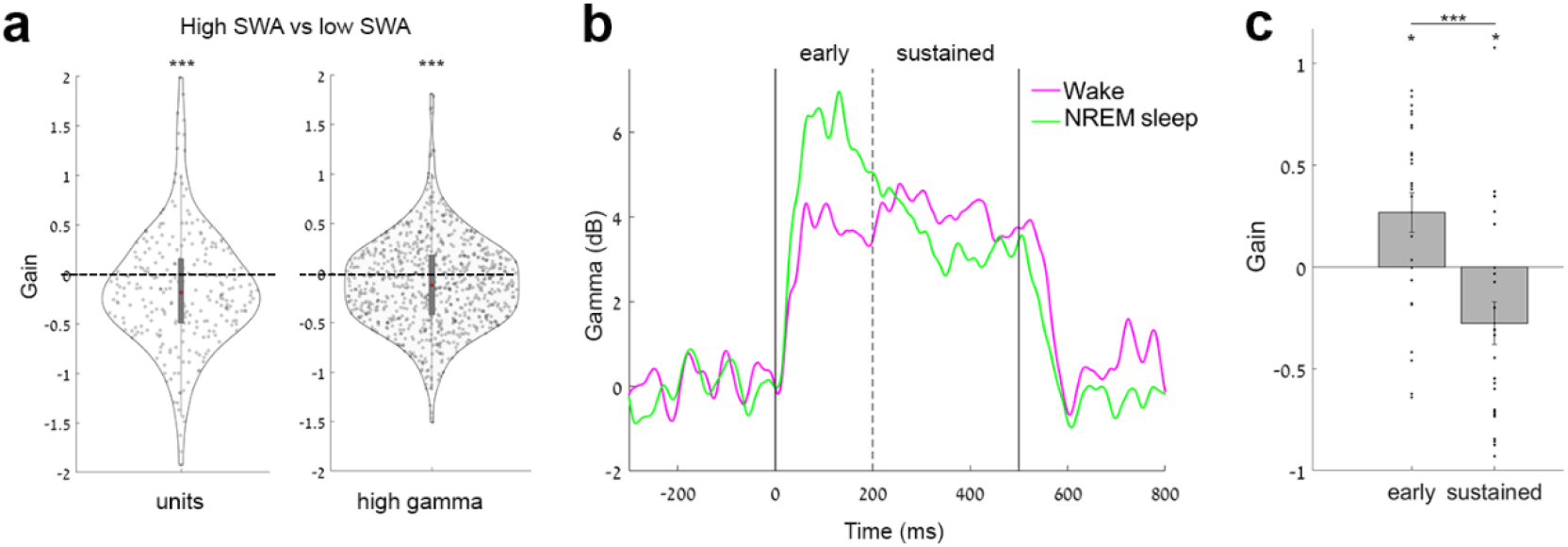
Factors associated with the degree of auditory response attenuation in NREM sleep. **(a)** (left) Auditory spike responses during NREM sleep with high (top 20%) SWA show stronger attenuation (negative gain on y-axis) compared to periods of low (bottom 20%) SWA (n = 263 spike responses from 55 clusters; attenuation of −16.10%, p = 5.9*10^−8^ by signed-rank test). (right) same association between high SWA and greater sleep attenuation for auditory high-gamma LFP responses (n = 527 high-gamma LFP responses from 74 microwires; attenuation of −10.73%, p = 6.8*10^−05^ by signed-rank test). A similar effect was found for periods of high sigma (10-16 Hz) power representing spindle activities (gain = −7.04%, p = 3.9*10^−06^ for high gamma; gain = −8.31%, p = 5.3*10^−06^ for spiking, signed-rank test). **(b)** Representative low-gamma response to a 40 Hz click-train in wakefulness (pink) and NREM sleep (green) shows differences between early vs. sustained response components. **(c)** Quantitative analysis across all low-gamma responses to 40 Hz click-trains (n = 25 microwires) reveals that sustained responses show a stronger attenuation than early response during NREM sleep (p = 5.1*10^−05^ by signed-rank test). Early responses were actually slightly potentiated during NREM sleep (positive gain of 26.65%, p = 0.016 via signed-rank test).

**Extended Data Fig. 7.**
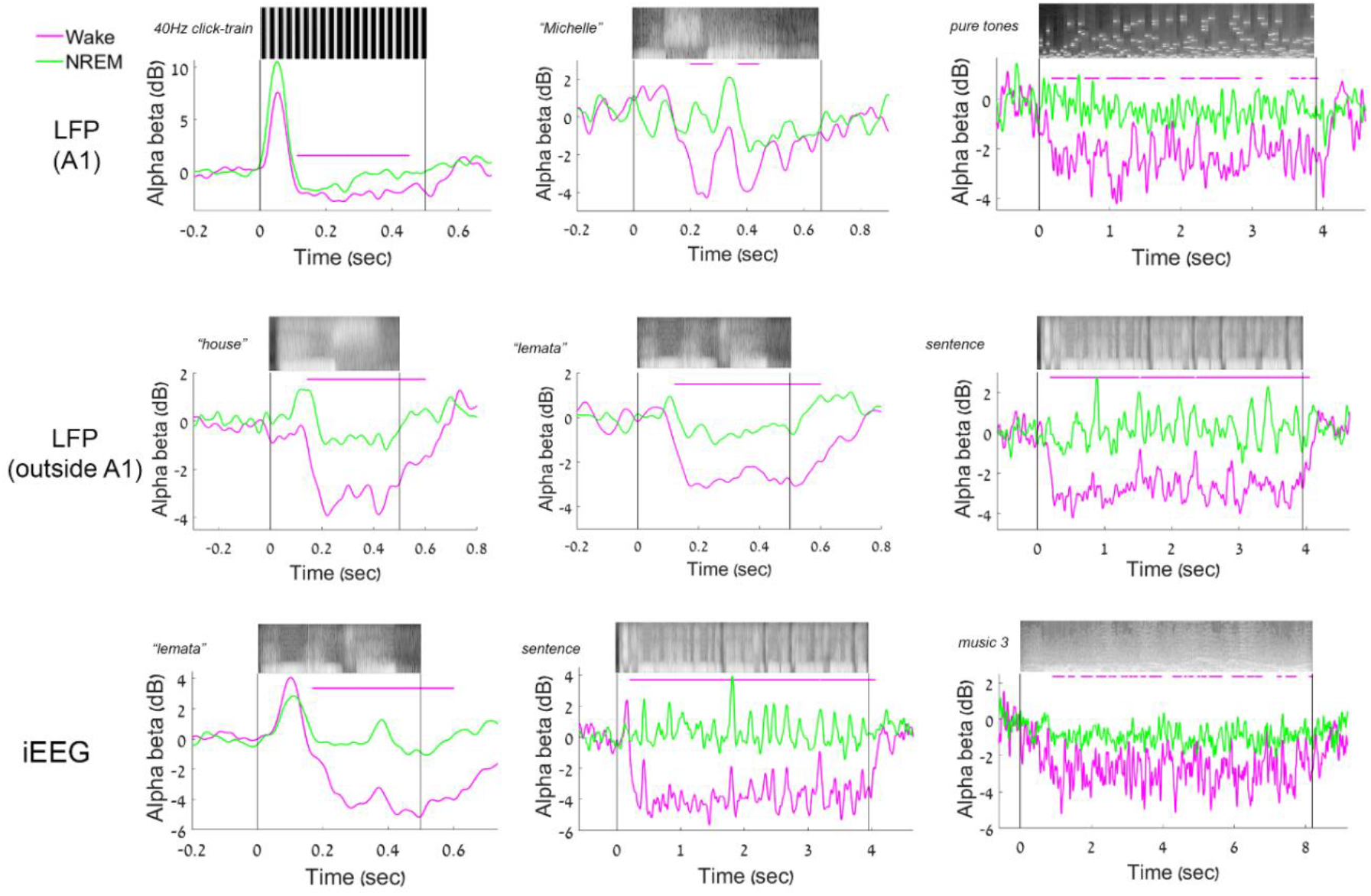
Additional examples of LFP alpha-beta auditory responses. LFP induced alpha-beta (10-30 Hz) power time-courses during wakefulness (pink) and NREM sleep (green) reveal disrupted alpha-beta responses during sleep.

**Extended Data Fig. 8.**
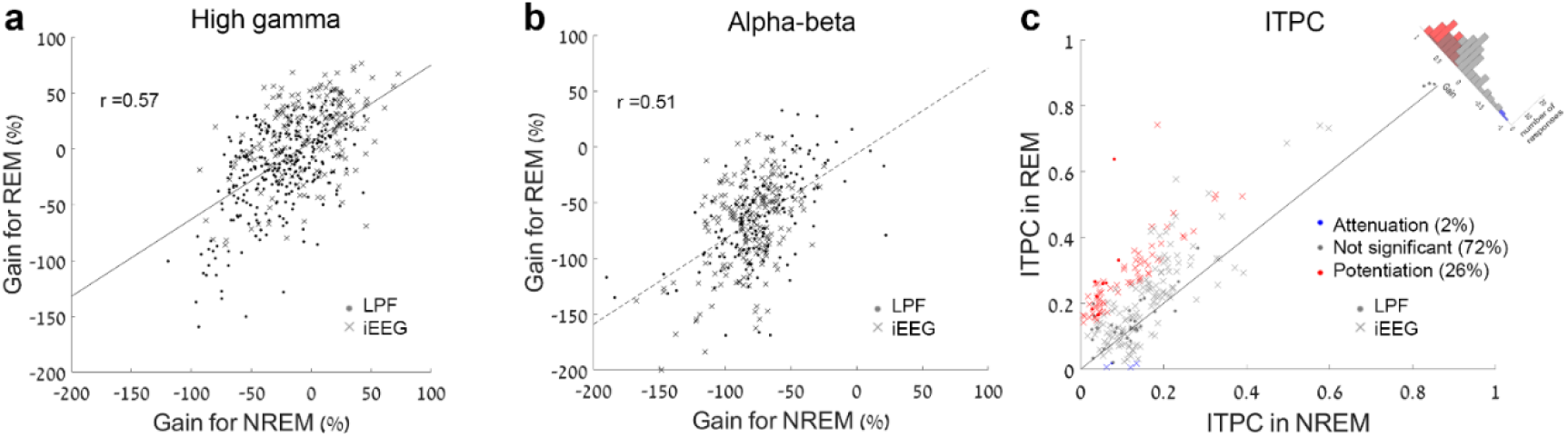
Comparing responses in NREM and REM sleep. **(a)** Scatter plot comparing the degree of sleep attenuation for high gamma power auditory responses (n = 89 responses from LFP microwires and 86 responses from 15 iEEG electrodes) in REM sleep (y-axis) vs. NREM sleep (x-axis) shows significant correlation (r = 0.72, p < 0.001). **(b)** Same as (a) for ABD (n = 111 responses from 16 LFP microwires and 92 responses from 11 iEEG electrodes, r = 0.72, p < 0.001). **(c)** Scatter plot of inter-trial phase coherence (ITPC) of 40 Hz iEEG and LFP responses (n = 43 and 202 respectively) in REM sleep (y-axis) vs. NREM sleep (x-axis), together with a histogram of gain values comparing response magnitudes (along the diagonal). The results reveal partial potentiation of the responses during REM sleep compared to NREM sleep (26% stronger entrainment, mean gain: +17.20%).

